# Quantifying internal conflicts and their threats to organismal form and fitness

**DOI:** 10.1101/2024.02.05.578856

**Authors:** Martijn A. Schenkel, Manus M. Patten, J. Arvid Ågren

## Abstract

Evolutionary biologists often treat organisms as both fitness-maximizing agents and as the primary level at which adaptation manifests. Yet, genes and cells may also seek to optimize their fitness by distorting the Mendelian rules of transmission or by influencing organismal traits for their own benefit. Organismal form and fitness are therefore threatened from within by selfish genes and cells. However, to what extent such internal conflicts actually harm individual organisms and threaten our concept of the organism as the sole bearer of adaptation remains unclear. We introduce a mathematical framework to capture the threat posed by internal conflicts and develop two metrics to measure their various forms of harm. We name these metrics fitness unity and trait unity, and use them to refer to the threats posed by internal conflicts to an organism’s role as the optimizing agent and the strategy wielded to achieve that optimization, respectively. We apply our framework to two examples of internal conflicts, genomic imprinting and sex ratio distortion, to illustrate how such harms from internal conflict may be quantified. We conclude by discussing the conditions under which internal conflict becomes sufficiently disruptive to organisms that it no longer makes sense to think of them as unified fitness-maximizing agents, but instead as adaptive compromises of multiple competing sub-agents.

## Introduction

Understanding the apparent design of living organisms is one of the major goals of evolutionary biology. To that end, organisms can be modelled as optimizing agents that strive, and sometimes manage, to maximize their inclusive fitness (Grafen, 2006, 2014; Levin & Grafen, 2019). This framework to adaptation has been especially popular and productive in its applications in behavioral ecology and social evolution more generally (Davies *et al*., 2012; West & Gardner, 2013). Associated with this approach is the view that organisms are the main bearers of adaptations. That is, if natural selection is conceived as a would-be designer of sorts (e.g., Lewens, 2004; Dennett, 2009), then its designs (adaptations) are for the benefit of the individual organism (Gardner, 2009). The existence of selfish genetic elements and selfish cell lineages, however, threatens both of these views (Ågren, 2016; Gardner & Úbeda, 2017; Okasha, 2018; Scott & West, 2019; Howe *et al*., 2022, 2024). Such internal conflict can prevent organisms from maximizing their inclusive fitness, and it leaves open the possibility that adaptive phenotypes can be found at sub-organismal levels of selection.

Recently, we have outlined how selfish sub-organismal elements use two main strategies to achieve this: distorting their numerical representation (transmission distorters) or distorting organismal traits in order to maximize their own fitness rather than that of their organism (trait distorters) (Patten *et al*., 2023). A quintessential example of a transmission distorter is a homing endonuclease gene. This is a stretch of DNA that, when present in a heterozygous state, causes double-strand breaks in the alternative allele and then uses itself as a template for repair, overwriting the alternative allele with its own sequence. Through this process, the organism goes from heterozygote to homozygote, which guarantees that the homing endonuclease is transmitted to the next generation, with little to no cost to organismal fitness (Burt & Koufopanou, 2004; Oberhofer *et al*., 2018). In contrast, trait distorters do not alter the mechanics of transmission – they follow Mendel’s laws of fair transmission – but alter an organismal phenotype in some way to enhance their own fitness. A prime example of this strategy is that of genomic imprinting, where the expression of an allele depends on whether it was inherited from the mother or father (Moore & Haig, 1991; Haig, 2000), and the organism’s phenotype has different inclusive fitness consequences for the maternally- and paternally-derived alleles. Pure transmission distortion does not necessarily harm organismal fitness—at least, not in theory—but often transmission distorters are associated with side effects that could be classified as trait distortion (e.g., disruption of spermatogenesis in some post-meiotic killers). Transmission and trait distorters are widespread across the tree of life (Burt & Trivers, 2006; Ågren & Clark, 2018), but opinions differ over their implications. On the one hand, they are viewed as fundamental to understanding genetics and as a source of evolutionary innovation (e.g., “…the prevalence of genomic conflict is universal, and it influences all aspects of genetic form and function”; Rice, 2013), but on the other hand, they have been brushed aside as mostly inconsequential for organismal adaptation (e.g., “The disadvantages of ignoring genetic conflict will often be minor, because we expect it to have relatively little influence on adaptation at the individual level”; West & Gardner, 2013). This disagreement points to the need for a formal measure of their influence.

In this paper, we present a mathematical framework to quantify how internal conflicts undermine organismal adaptation. Our approach allows us to distinguish two measures of organismal disunity, separately addressing whether the harm from internal conflict has to do with traits or fitness. To illustrate the value of the framework, we apply it to a pair of internal conflicts where organismal traits are subject to selection at both the organismal and sub-organismal level: genomic imprinting and sex ratio distortion. We conclude by discussing whether internal conflicts may ever be sufficiently potent as to warrant a reconsideration of organisms as the only true agents in the evolutionary process and the lone bearers of adaptation.

### A mathematical framework to quantify internal conflicts

To capture the conflict between organismal and sub-organismal entities we use the mathematics of optimization, conceiving both genes and individuals as fitness-maximizing agents (Grafen, 1999; Gardner & Welch, 2011). While our focus is here on conflicts between genes and individuals, the framework can readily be adapted to conflicts between cells and individuals. We first present a general framework before applying it to some specific scenarios below. Unless otherwise specified, the specific models are identical to this general model (see Table 1 for an overview of all variables and parameters in both the general and specific models). Throughout, we use subscripts G and I to refer to gene- and individual-level variables and functions, respectively. Individual fitness is given by the function *w*_I_(*z*_I_) where *z*_I_ is an individual’s phenotypic trait subject to selection. This establishes an individual-level optimization program such that:

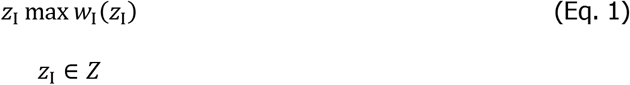

**Table 1:**
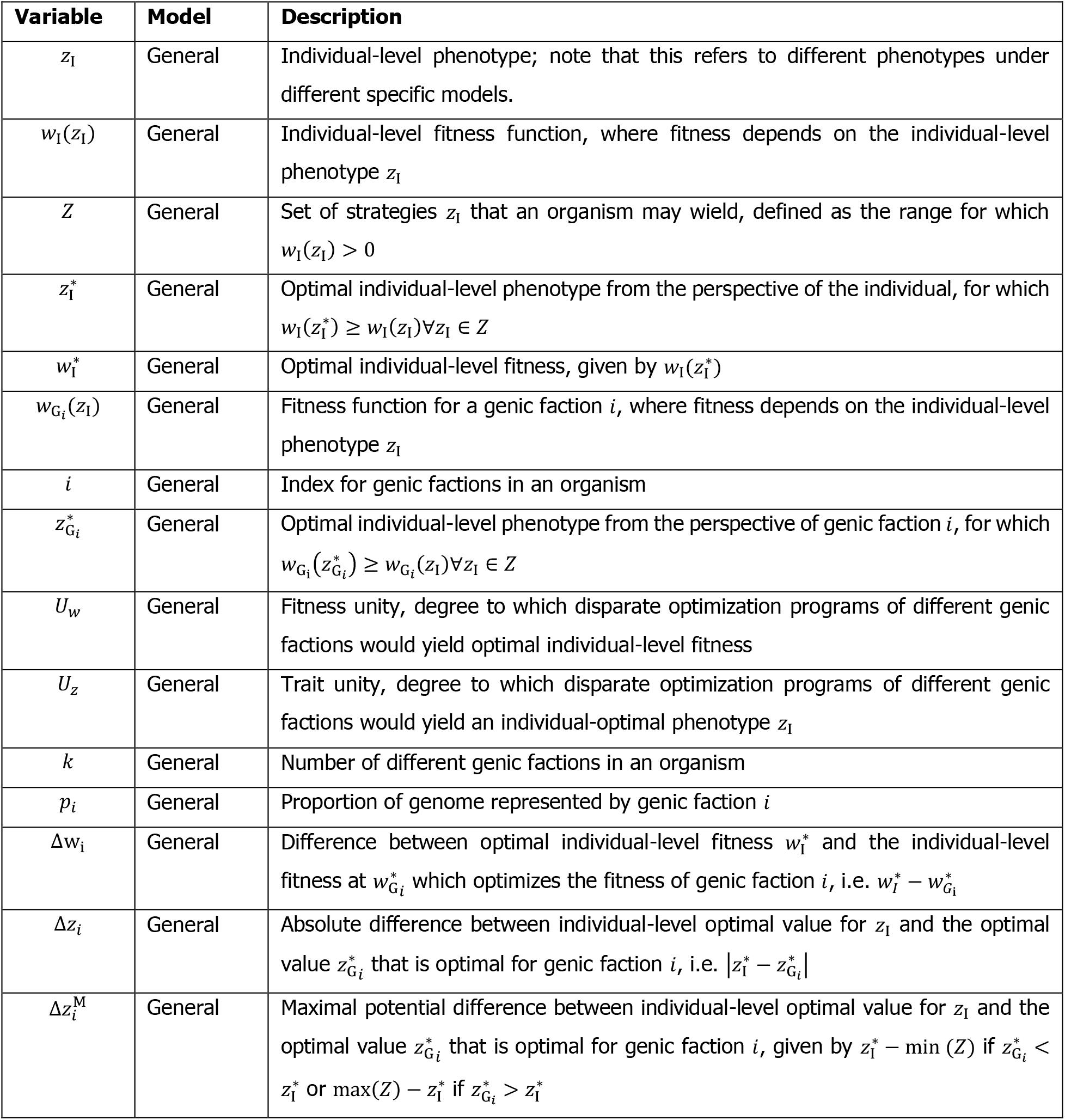
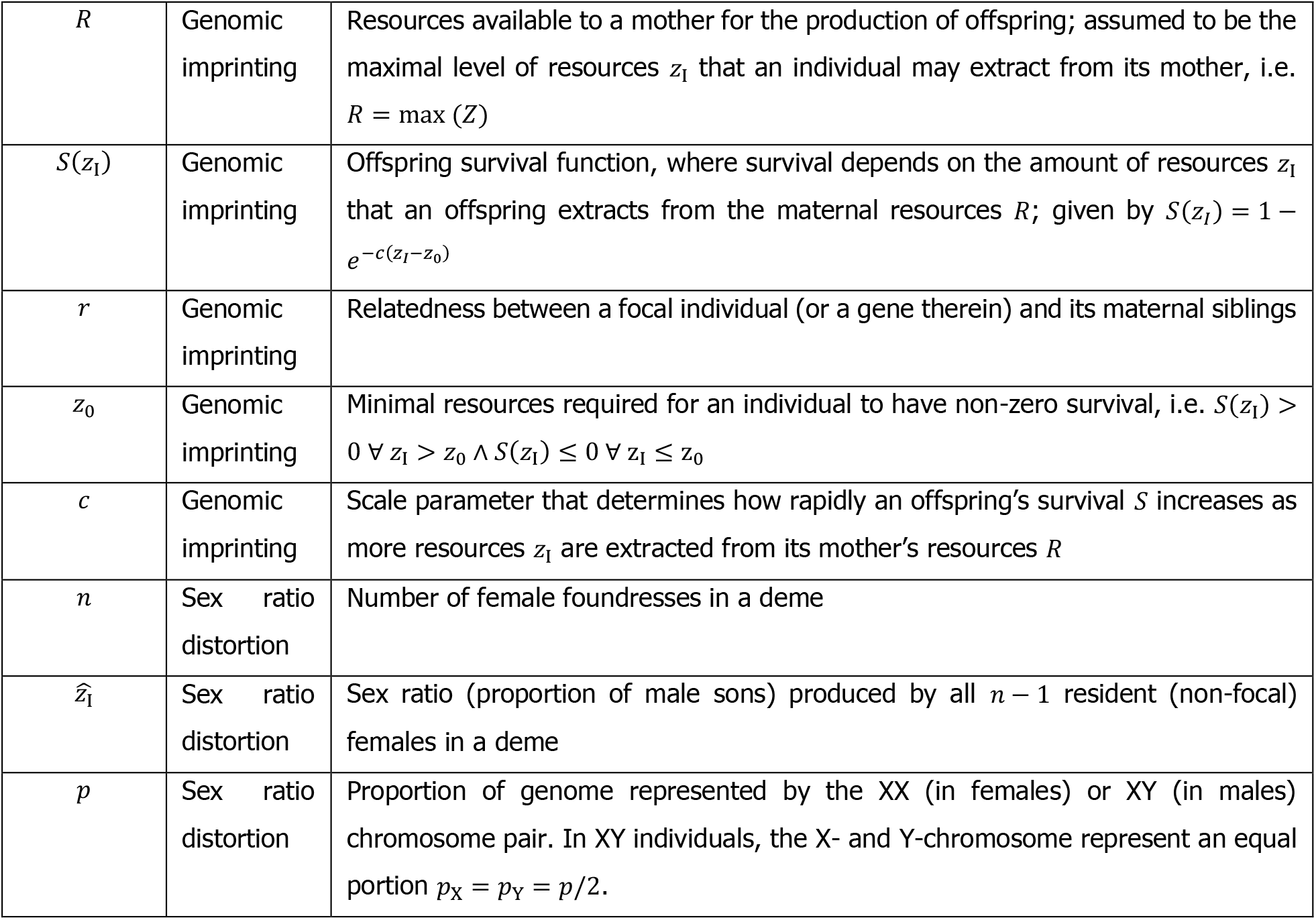
Definitions and usage of all model variables and functions.

The value of *z*_I_ is selected from a set of possible trait values *Z* so that the value of *w*_I_(*z*_I_) is maximized; we denote this value as 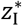, and the resulting optimal fitness 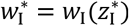. Individual fitness *w*_I_ is taken to be equivalent to an individual’s inclusive fitness. Genes, being located within organisms, are subject to selection based on the individual-level trait in a similar way. We assume that each gene *i* has a fitness 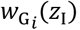, and accordingly an optimization program given by:

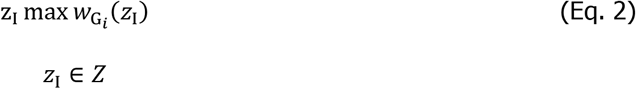

Hereafter, we refer to genes that share the same optimization program as belonging to a specific genic faction. A genic faction is similar to the coreplicon concept defined by Cosmides and Tooby (1981). However, whereas their concept is primarily focused on the transmission pattern of different genes, our classification differs in that we also consider other subdivisions of the genome, such as by parental origin (maternal versus paternal), to allow for other factors that may lead to different optimization programs between factions (c.f. Gardner and Úbeda 2017). For each genic faction *i* ∈ {1,2, …, *k*} there exists a value of *z*_I_ ∈ *Z*, denoted 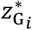 that maximizes 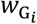. Genic fitness 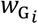 is defined as the inclusive fitness of a gene in faction *i*, the number of gene copies that are transmitted to the next generation by either the focal organism or its relatives, or similarly the degree of genetic representation at an undefined future time. Because we define our genic factions according to specific optimization programs, the inclusive fitness of a single gene in this faction should hold for all genes in that faction.

We present two metrics, *U*_*z*_ and *U*_*w*_, which depend on the extent to which internal conflicts affect individual trait values (*U*_*z*_) or individual fitness (*U*_*w*_), respectively (Figure 1). We christen these measures ‘trait unity’ and ‘fitness unity’. Verbally, fitness unity can be thought of as the extent to which natural selection would result in individual fitness optimization if free rein were given to genic factions. We consider the optimal outcome for each faction separately, and weigh these according to their share of the individual’s genome. In other words, it describes how well the individual optimization program would be represented by the disparate optimization programs of its constituent parts. Trait unity, in contrast, captures the extent to which trait values would reflect the organismal optimum under the same assumptions. It describes how aligned the constituent parts are in their construction of a specific organismal phenotype.

**Figure 1:**
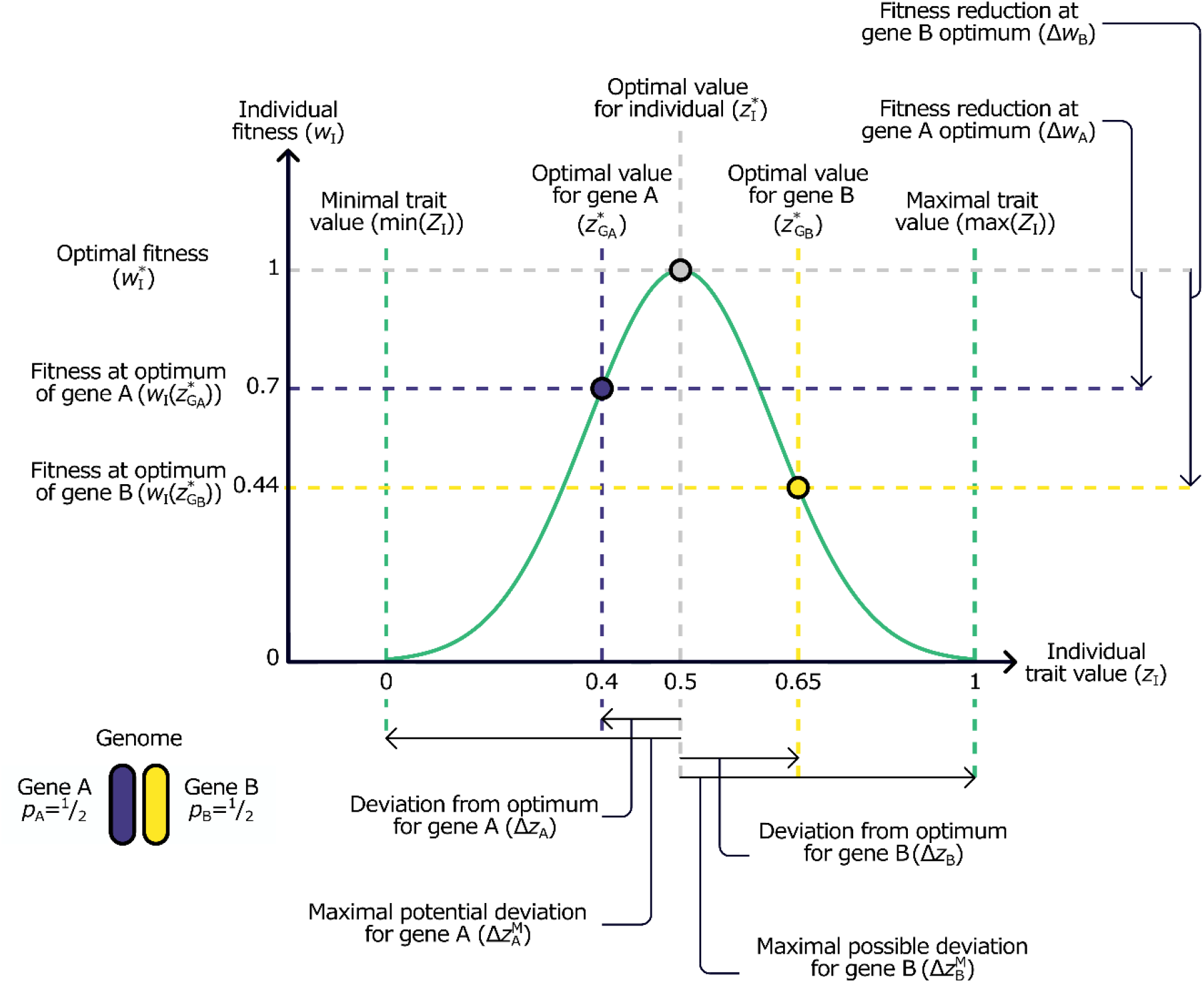
Individual fitness, individual versus genic faction optima, and organismal unity. Individuals are represented by a simple genome consisting of just two genic factions, *i* ∈ {A, B}, shown in blue and yellow. We use “genic faction” to refer to a collection of genes that share an optimization program, which may or may not be identical to the individual-level optimization program. Individual fitness *w*_I_ (green curve) is determined by a trait *z*_I_ ∈ *Z* for which *w*_I_(*z*_I_) > 0, and is optimized for 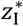 (grey dot and lines); genes A and B optimize their fitness when *z*_I_ equals 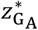 (blue dot and lines) and 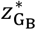 (yellow dot and lines). Arrows below the graph indicate the difference Δ*z*_*i*_ between the individual-optimal value 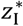 and the genic optima 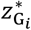, as well as the maximum difference 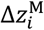 depending on whether a genic optimum is lower than the individual-level optimum 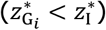 or vice versa. Arrows to the right of the graph indicate the difference between the individual-optimal fitness score 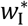 and the individual fitness at genic optima 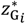.

Under these assumptions, fitness unity (*U*_*w*_) and trait unity (*U*_*z*_) are given by:

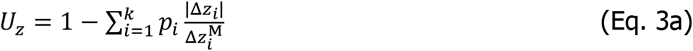

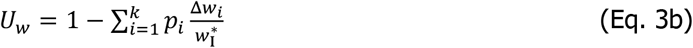

In Eqs. 3a,b, *p*_*i*_ represents the share of the total fitness or phenotype attributable to genic faction *i* (so that *∑*_*i*_ *p*_*i*_ = 1). The definition of *p*_*i*_ is intentionally flexible and we interpret it here as a measure of how much of the genome follows a specific optimization program. As such, it can mean the number of genes, linkage groups, or even base pairs, depending on how one carves up the genome. The capacity of a genic faction to determine a contested trait could be a specific, empirically measurable quantity, like the faction’s current influence on the trait. Alternatively, it could reflect some reasonable evolutionary potential to affect the trait, which may vary because only some factions may hold specific genes that map onto the phenotype of interest. We explore these different interpretations of *p*_*i*_ in the Discussion. In Eq. 3a, Δ*z*_*i*_ reflects the distortion of the trait value *z*_I_ away from 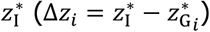. We divide the absolute value of this difference by 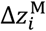, which is the maximal potential distortion between the individual-optimal value 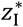 and 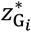. Its value, depending on whether 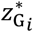 exceeds 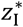 or vice versa, is given by:

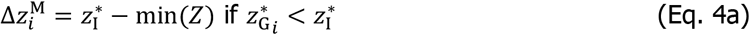

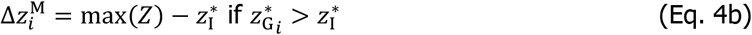

The difference between 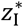 and each genic faction’s optimum 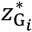 is most severe when 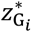 equals the minimum (when 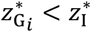) or maximum value (when 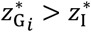) of *Z*, where *Z* denotes the range of values for *z*_I_ where *w*_I_(*z*_I_) > 0, or alternatively a similarly constrained range of values, e.g. [0,1] when traits represent rates or proportions. In Eq. 3b, Δ*w*_*i*_ represents the difference between an organism’s optimal fitness 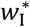 and its fitness at the trait value 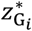 that is optimal for gene *i*, 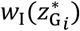 where 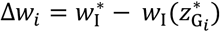. We divide this difference by 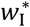 to obtain the relative reduction in individual fitness. A detailed example to illustrate how these calculations work is provided in Box 1.

Our measurements of organismal unity function as kinds of counterfactuals. Both are about the extent to which theoretical optima are aligned—not the extent to which genes and/or organisms achieve these optima. They do not reflect whether organisms are actually fitness-optimal, or whether certain genic factions are actually successful in distorting organismal traits. Instead, the framework quantifies the extent to which the genic factions in an organism *could* harm organismal fitness or *could* disagree over what would be optimal. In other words, it answers the following question: to what extent do the various genic optimization programs align with the individual optimization program? Just because an element cannot actually distort the trait away from the organismal optimum, it does not mean that it optimizes its own fitness at the organismal optimum.

### Box 1: Calculating trait unity and fitness unity

To illustrate how trait and fitness unity can be calculated, we consider the scenario presented in Figure 1, where individuals have a positive fitness score (green curve) when 0 < *z*_I_ < 1.0, and their fitness is maximized at 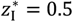 (grey dot and lines). Each individual carries two genic factions A (blue) and B (yellow) that both represent one half of the genome 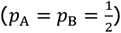. These factions have optimal trait values of 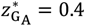 and 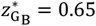 (indicated by the blue and yellow dots and lines for genes A and B, respectively). The absolute differences between the faction optima and the individual optimum then are Δ*z*_A_ = |0.5 − 0.4| = 0.1 for faction A and Δ*z*_B_ = |0.5 − 0.65| = 0.15 for faction B. Because 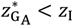, the maximal potential distortion for faction A is given by 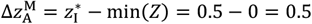. Inversely, because 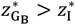, we find that 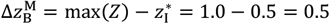 for faction B. We thus obtain for trait unity *U*_*z*_:

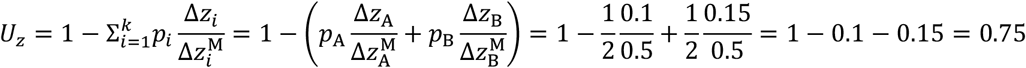

To determine the fitness unity, we need to consider what individual-level fitness would be attained at the optima for genic factions A and B. In our example, individual fitness is optimized for 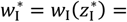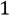. Plugging in the genic optima for two factions, we find that 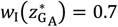, and 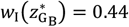. The differences between the individual optimum and the genic optima are then Δ*w*_A_ = 1 − 0.7 = 0.3 and Δ*w*_B_ = 1 − 0.44 = 0.56. We then obtain for fitness unity *U*_*w*_:

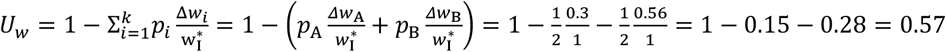

Table 2 provides a detailed overview of the above calculations for trait and fitness unity.

**Table 2:**
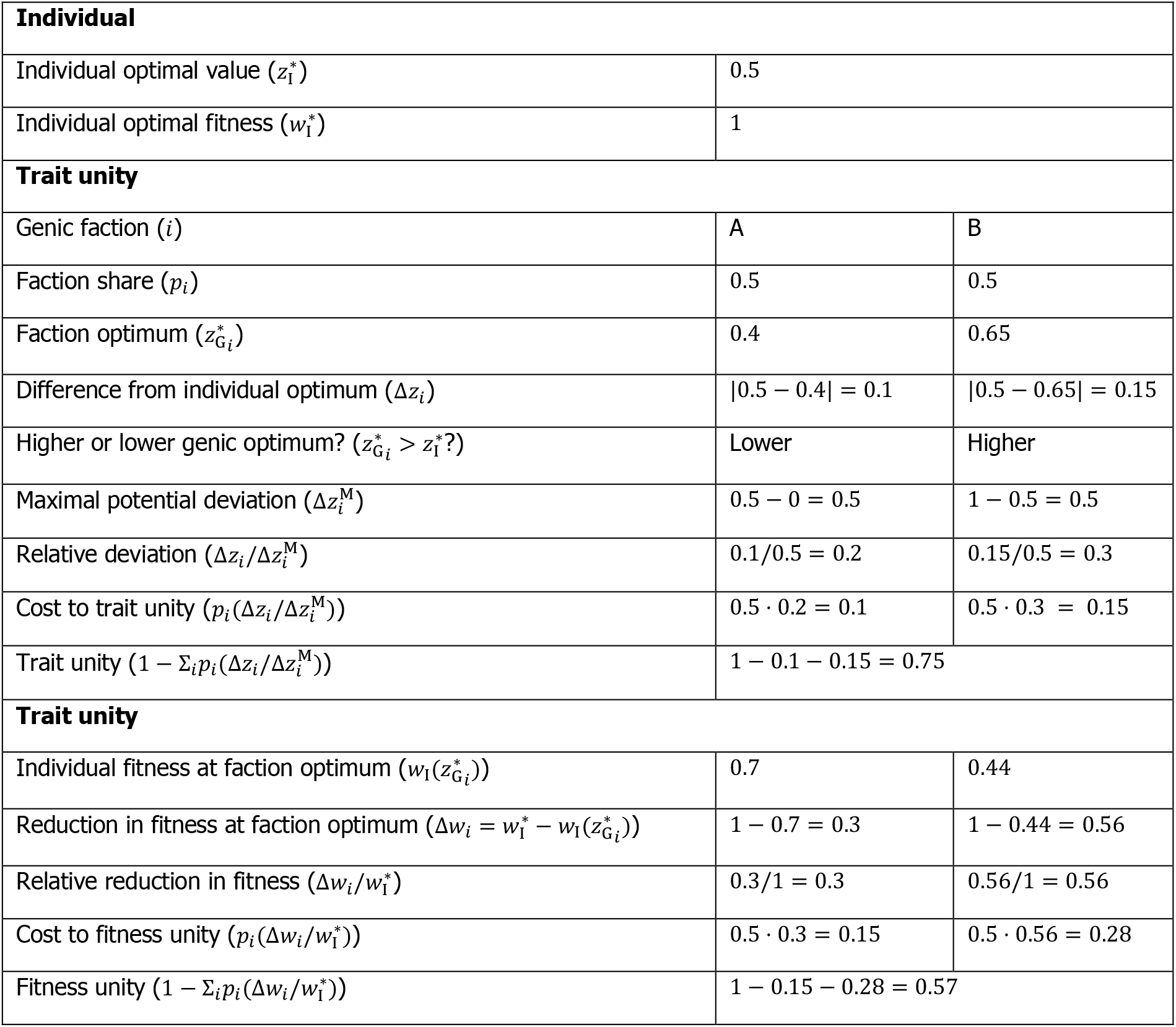
Example calculations for trait and fitness unity.

### Applying the framework: Modelling specific examples of internal conflicts

To illustrate our framework with something more concrete, we explore two examples of internal conflicts: genomic imprinting and sex ratio distortion. Unless otherwise specified, both models assume a large (possibly infinite) panmictic sexually-reproducing population consisting of diploid individuals. Mating yields juvenile offspring and the production of sons and daughters is equally costly. All juveniles have equal survival to adulthood. Generations are non-overlapping, as the cohort of matured juveniles replaces the previous cohort of adults. In both the imprinting and sex ratio distortion models, our goal is to illustrate how organismal disunity may be quantified, not to model all aspects of the specific biology, and simplifying assumptions are made accordingly. Further details for both models are provided in the Supplementary Material.

#### Genomic imprinting

Imprinted genes are expressed at different levels depending on whether they were inherited maternally or paternally. Haig’s kinship theory posits that genomic imprinting evolved because the maternally- and paternally-derived genome sets (matrigenes and patrigenes) experience different relatedness to the bearer’s kin (Haig, 2000; Patten *et al*., 2014), which will affect the inclusive fitness of the two genomes differently. It is this divergence of interests that we model here. In the Patten *et al*. (2023) classification, imprinted genes are trait distorters that alter the traits of organisms, but are transmitted in a fair, Mendelian manner. Our model assumes a mother who has *R* resources to be used to produce offspring, each of which is sired by a different male such that they are maternal half-siblings. An offspring’s survival *S* depends on the level of investment as given by a monotonically increasing function of the form:

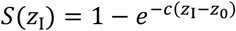

Here, *z*_0_ is the minimal investment required for producing viable offspring and *c* is a scaling parameter that controls how strongly survival increases with increasing investment. The expected number of surviving offspring produced by a mother is thus given by *RS*(*z*_I_)/*z*_I_. Here, relatedness refers to the probability that an allele sampled from the organism and an allele sampled from its kin are identical by descent. The matrigenic genome therefore has a relatedness 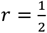 to any of the mother’s offspring, whereas for the patrigenic genome the relatedness is 0. The organismal relatedness is given by their average, i.e. *r* = 1/4. Maternal investment is assumed to determine offspring survival, as well as how many offspring a mother is expected to produce. Due to relatedness differences between the matrigenic and patrigenic genome halves and the offspring as an integral unit, the inclusive fitness benefit of a focal offspring’s siblings will be evaluated differently by each party. This difference causes each of them to experience different optimal levels of investment *z*_I_, which are evaluated to determine the organismal unity of purpose.

Under this model, the patrigenic and matrigenic genome of an offspring have different optimal levels of maternal investment, each of which differs from the organismal optimum 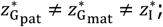 Supplementary Figure 1). The patrigenic genome, not being represented in the offspring’s kin, favors the investment level that maximizes offspring survival, which occurs when *z*_I_ = *R* so that 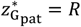. In contrast, the optimal investment level of the offspring does incorporate indirect fitness through kin survival, but these benefits are weighed at a relatedness 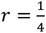. The matrigenic genome considers a similar tradeoff, but assumes a relatedness of 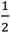. This causes the matrigenic genome optimum 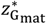 to be lower than the organismal optimum 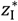, as the matrigenic genome more highly values the inclusive fitness benefit derived from kin survival.

The different interests of the matrigenic and patrigenic genomes both contribute to reduced trait and fitness unity (Figure 2; Supplementary Figures 2 and 3). However, their effects are substantially different. The matrigenic genome’s optimum value would result in lower organismal optimality when offspring survival increases less rapidly with increasing investment (low *c*) and/or maternal resources are scarce (low *R*; Supplementary Figure 2, left column). The patrigenic genome optimum value however is more harmful when resources are abundant (high *R*) and/or survival is relatively easy (high *c*; Supplementary Figure 2, right column). This reflects the matrigenic genome adopting a more favorable view of kin; this leads to reduced survival of the focal organism, while enhancing that of its kin. The patrigenic genome does the opposite; it enhances organismal survival, but completely disregards the survival of its kin. In terms of unity of purpose, we see that the interests of the patrigenic genome tend to dominate; both trait and fitness unity are most strongly reduced when resources are abundant (high *R*), particularly when return-on-investment *c* is high and/or minimal investment *z*_0_ is low (Figure 2).

**Figure 2:**
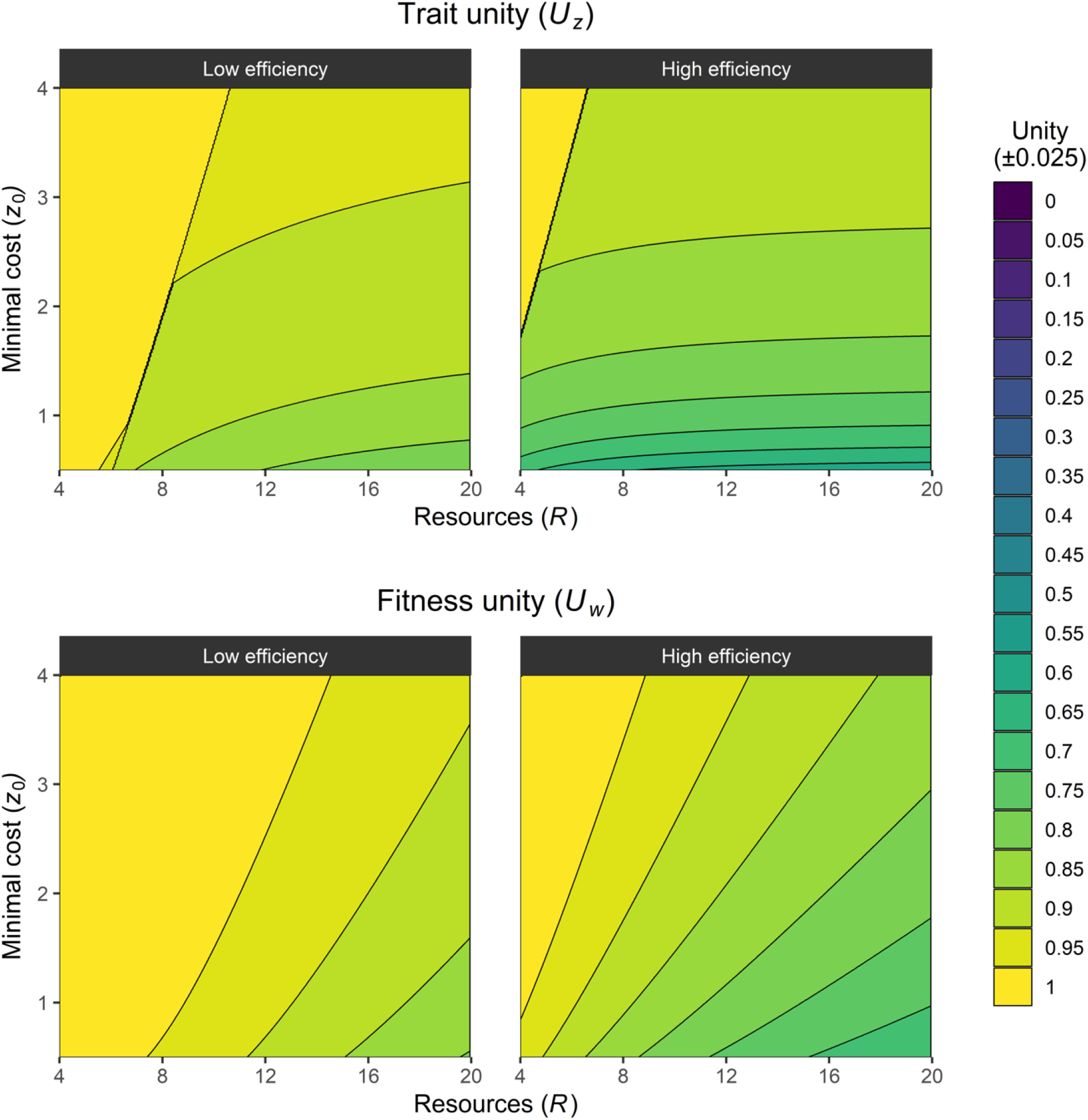
Trait unity (*U*_*z*_) and fitness unity (*U*_*w*_) of parental investment in light of genomic imprinting. Optimal parental investment is investigated from the perspective of their offspring; increased investment enhances their survival, but reduces the production of siblings. Owing to relatedness differences, matrigenes and patrigenes favor under- and overinvestment of resources relative to the individual-level optimum, reducing the trait unity. In particular, patrigenes favor complete investment of all maternal resources *R* as they have no representation in any of the other offspring produced. This difference drives reduced trait and fitness unity as it harms the indirect fitness component of the focal offspring’s inclusive fitness. *z*_0_ represents the minimal investment, whereas low versus high efficiency refers to how rapidly survival increases with further investment (low efficiency: *c* = 0.5; high efficiency: *c* = 1.5). Lower minimal investment (*z*_0_) and/or higher efficiency (*c*) leads to even stronger reductions in trait and fitness unity through the patrigenes’ preference for overinvestment.

### Sex ratio distortion

Sex determination is an essential component of individual development and successful execution of this process is essential for individual fertility and mate compatibility. Despite its importance, the genetic systems that govern this process are surprisingly variable and exhibit high evolutionary rates (Bachtrog *et al*., 2014; Beukeboom & Perrin, 2014). One key contributor to this divergence is genetic conflict (Werren & Beukeboom, 1998; Schenkel et al., under review). Different genetic elements within an individual segregate to and from males and females in different ways (Haig *et al*., 2014); autosomes move freely from parents to offspring of both sexes, whereas Y-chromosomes segregate exclusively from fathers to sons, and cytoplasmic material inherits almost exclusively through the maternal route (but see Munasinghe & Ågren, 2023). Genetic elements that are transmitted at unequal rates to and from individuals will prefer to bias sex determination towards that sex through which they are transmitted more often (Cosmides & Tooby, 1981; Werren & Beukeboom, 1998). By extension, these genetic elements may also be in conflict with their bearer over the optimal sex ratio among its offspring (Shuker *et al*., 2009) and reproductive parasitism by endosymbionts represents a prime example of how cytoplasmic elements may seek to distort sex ratios (Werren *et al*., 2008).

The evolutionary forces governing offspring sex ratio (defined here as the proportion of males produced among an individual’s offspring) have been extensively investigated (reviewed in West, 2009), which means the trait optima needed for the computation of our unity measures are already established. For example, in large populations, theory predicts that, all else equal, it would be optimal for individuals to produce an equal proportion of sons and daughters (Fisher, 1930). In structured populations, however, where individuals are distributed across small patches and interact with a limited number of individuals, the optimal sex ratio may deviate considerably from 1:1. Hamilton (1967) introduced the concept of local mate competition, where broods of offspring mate—or alternatively compete for mating—with their patch mates. Specifically, Hamilton considered a case where females mate locally and then disperse to found new demes, potentially alongside other females. Males mate with females in their natal patch only, but do not disperse. Under these conditions, an individual’s fitness is best represented by the number of grandoffspring that it produces, which is proportional to the number of daughters produced plus the number of females inseminated by its sons. We adopt a similar modeling approach to quantify organismal unity with regard to offspring sex ratio and consider how sex-biased transmission of the X- and Y-chromosomes result in deviations in optimal sex ratio for these genetic elements.

Optimal sex ratios for organisms are determined by the maximization of the number of grandoffspring. At the genic level, this corresponds to a general cumulative contribution to the gene pool in the second generation after the focal individual (i.e., F_0_ to F_2_). For the X- and Y-chromosomes, this criterion is adapted to reflect the number of X-as opposed to Y-chromosomes that are transmitted to this generation (Hamilton, 1979). In XX individuals, X-chromosomes favor a slightly more female-biased sex ratio (Supplementary Figure 4). In small demes, this bias leads to a small reduction in trait unity (Figure 3, top left), though fitness unity is almost perfectly maintained (Figure 3, bottom left). In XY individuals, however, the case is different. X-chromosomes favor a stronger female bias, whereas Y-chromosomes prefer a much more male-biased sex ratio, reflecting their strict inheritance to females c.q. males. The observation of lower unity in the heterogametic sex is mirrored in a ZW system; Supplementary Figure 5. While at low deme sizes the differences between the individual optimum and the genic optima are not yet so severe, higher deme sizes yield optimal genic strategies that are approximately equal to all-female (X-chromosome) or all-male (Y-chromosome). Hence, trait unity at high deme sizes becomes maximally reduced, as both the X- and the Y-chromosome have genic optima that deviate maximally from the individual optimum (Figure 3, top right panel). In terms of fitness unity, however, the reduction is most apparent at intermediate deme sizes (Figure 3, bottom right panel). This is because at low deme sizes, the deviation in sex ratio is not very severe, whereas at high deme sizes, the reproductive values of sons and daughters come to be more or less equal. At intermediate deme sizes, the distortion in sex ratio, combined with the unequal reproductive values of sons and daughters, results in a stronger reduction in fitness unity than under other conditions.

**Figure 3:**
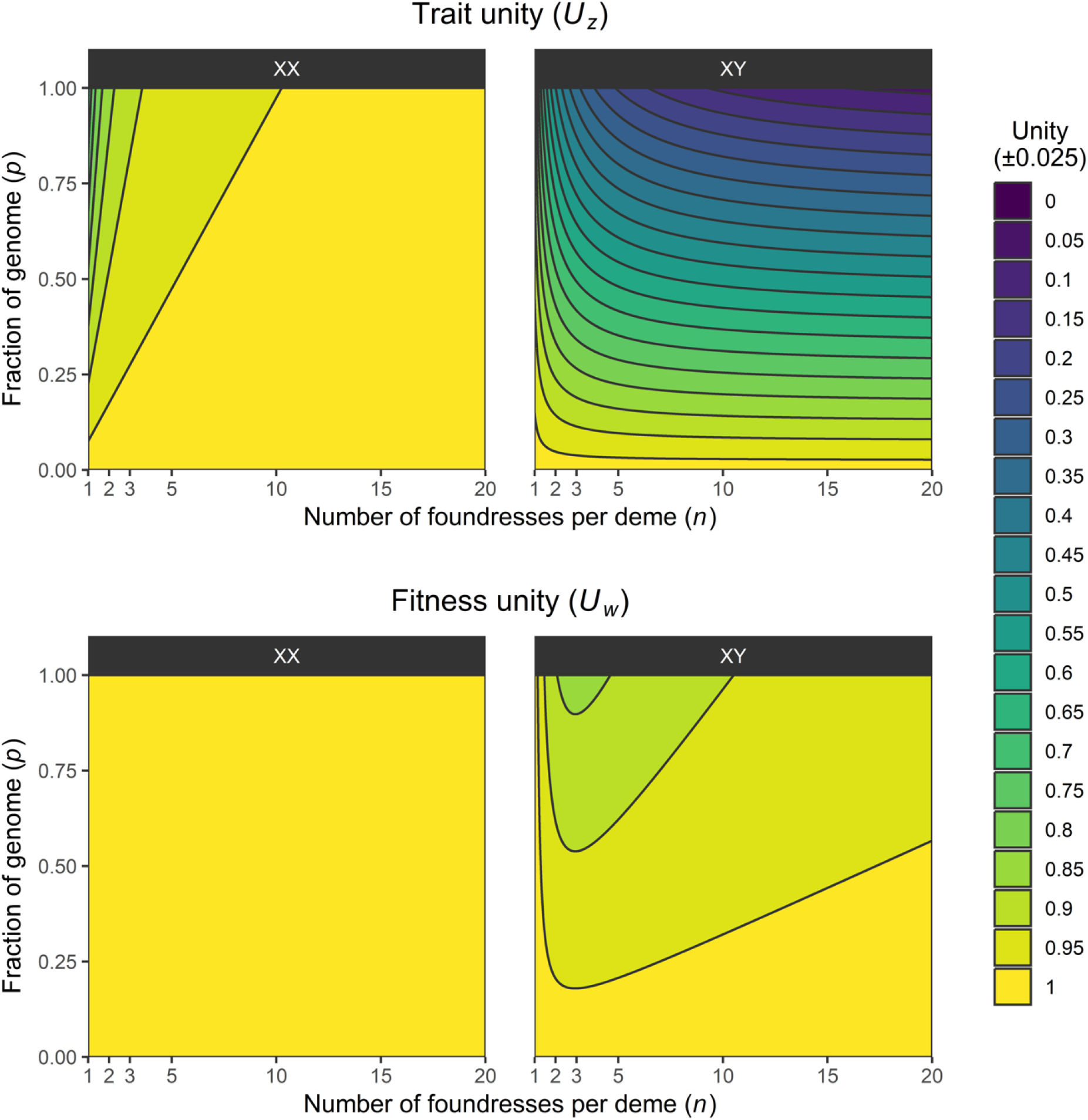
Trait unity (*U*_*z*_) and fitness unity (*U*_*w*_) of sex ratios under local mate competition in light of sex-biased inheritance. In determining trait and fitness unity, all *n* − 1 residents in the patch play the individual-optimal strategy 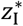. The focal individual consists of a genome of which a proportion 1 − *p* is autosomal, and therefore has the same optimal strategy as the individual. The remainder *p* is distributed among the X- and Y-chromosomes, which have genic-optimal strategies 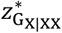 (in XX individuals), 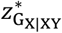, or 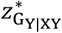 (in XY individuals); in XY individuals, the X- and Y-chromosome represent an equal portion *p*/2 of the genome.

## Discussion

Organismal adaptation is one of evolutionary biology’s foundational problems. Students of the issue have often treated organisms as both fitness-optimizing agents and the unique beneficiaries of adaptation. Both views are threatened by the existence of selfish genetic elements and selfish cell lineages that distort organismal form and fitness. Dawkins (1990) coined the term ‘The Paradox of the Organism’ for the observation that organisms persist, despite the ubiquitous potential for internal conflicts to tear them apart from within. Part of the resolution to this paradox seems to be that organisms can withstand a certain amount of internal conflict, a degree of internal disunity of purpose. Just how severely disunited organisms can be without breaking down altogether has been unclear. Furthermore, different organisms may harbor higher levels of disunity owing to their life cycle and developmental program (e.g. aggregative versus clonal multicellular organisms, haploid males versus diploid females in haplodiploids), which may differently affect their capacity for adaptation. To provide a foundation for resolving the paradox and formally studying organismal disunity, we introduce a mathematical framework to quantify the level of internal conflict that exists within organisms. Taking internal conflicts seriously raises several conceptual issues, including who or what is in control of the process of adaptation, what an organism is, and whether organisms can be undone from within. We address these issues in the following sections.

### Who is the organism? Or what is it that has an “organismal optimum”?

To quantify organismal disunity, we decompose the organism into a series of genic factions that experience different optimization programs, and so have different evolutionary optima. In this approach, each faction is scaled by its relative share, resulting in measures of trait and fitness unity that are effectively weighted means. These factional optima are then compared to the organism’s optimum. Ultimately, this is an organism-centric approach in that the organism’s optimum is taken as the reference, and the resulting unity measures describe how far the factions depart from this. But how should the organismal optimum be calculated in practice? One option is to take some part of the organism to represent the whole (c.f. Fromhage & Jennions, 2019). In practice, this usually means some unimprinted, eumendelian, autosomal reference gene’s optimum. Another is to use the weighted mean of the various genic factions’ optima. Different definitions may yield different values of 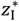, thus shifting the reference point for determining the overall measure (Figure 4).

**Figure 4:**
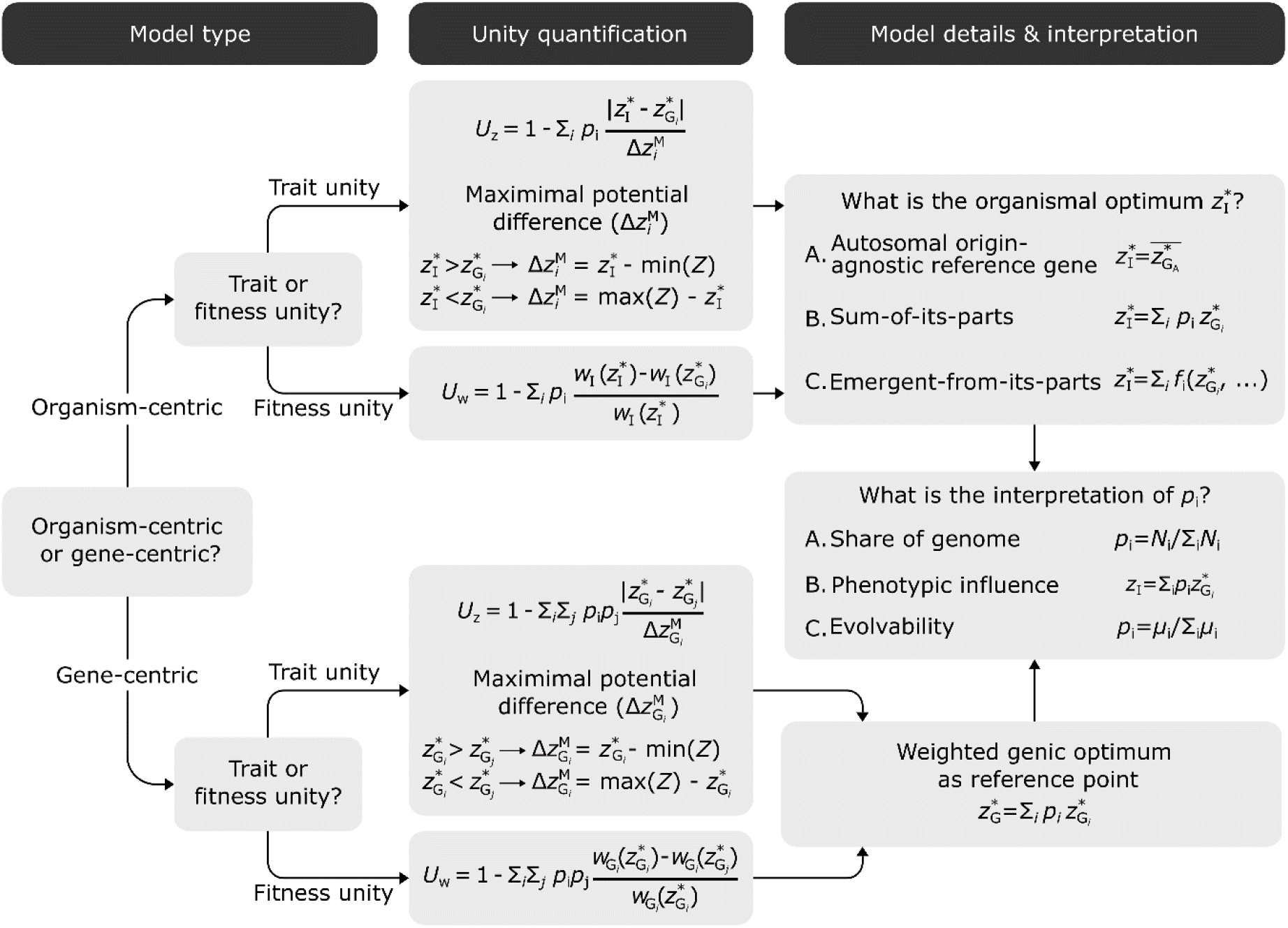
Alternative measures of organismal unity versus factional disunity. Different perspectives on internal conflicts within organisms can be developed by taking a gene-centric, rather than an organism-centric approach. Similarly, what constitutes the organismal optimum 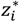 may be defined in several ways. Finally, a genic faction’s share *p*_*i*_ may be taken to mean different things to yield different classes of models. In our approach, we adopt an organism-centric approach (top branch) to quantify both organismal trait unity and fitness unity. In doing so, we assume that the organismal optimum is best represented as the optimum of an origin-agnostic autosomal reference gene, and that *p*_*i*_ represents the share of the genome represented by a genic faction *i*.

An alternative to an organism-centric approach is be to adopt a fully gene-centric approach and circumvent the organism altogether (Clarke, 2016). The gene’s-eye view sees organisms as temporary assemblies of conjoined genic factions that seek to exploit the resulting aggregate to further their own agendas (Ågren, 2021). Rather than determining deviations from a difficult-to-define organismal optimum, we could replace Eqs. 3a,b with something that uses the differences between optima for the different factions directly (Figure 4).

### What is a share of fitness or phenotype?

In our approach, *p*_*i*_ represents the share of the genome that can be assigned to genic faction *i*. The resulting measurements quantify the degree of disunity of an organism composed of a set of genic factions, each of which holds a certain share and follows a specific optimization program. As currently modelled, they are agnostic about whether these factions actually have the means to influence the phenotype under conflict, or how this capacity may evolve or affect evolutionary dynamics. Such agnosticism may become an issue. For example, the difference in transmission pattern between mitochondrial and nuclear genes places them in different genic factions, but the difference in gene number means that we expect the fitness interest of the latter to have a bigger influence over a disputed trait.

There are several ways these issues can be approached. One way would be to assign each genic faction with a certain power to influence the disputed trait value, so that the realized trait value is the aggregate of each faction’s optimal value, weighted by their power. Alternatively, we might interpret *p*_*i*_ as a measure of evolutionary potential, i.e. evolvability. Genic factions are likely to differ substantially not just in terms of number of genes, but also in mutation rates, regulatory complexity, etc. that may determine how rapidly they might be able to evolve in response to internal conflicts. Factions that have higher evolutionary potential should, over evolutionary time scales, be able to more frequently and/or more strongly distort organismal trait values to their benefit (similar to Scott & Queller, 2019; Rautiala & Gardner, 2023). By modeling organismal unity as a measure of evolutionary potential, this approach would describe the dynamics that are produced by specific internal conflicts.

Both approaches provide ways to go beyond our current approach which (merely) quantifies the latent severity of internal conflict. The price of the extension, however, is that they require additional, specific assumptions about the underlying biology. To determine the power or potential of a genic faction, we need to know which genes it contains that actually map to the disputed trait, and how this genotype-to-phenotype mapping depends on interactions with genes within its faction as well as in other factions. Understanding the evolutionary potential, and the ensuing dynamics, in turn requires in-depth knowledge of how these individual genes and genetic networks may evolve. Our framework provides a starting point for developing more detailed models on internal conflicts by establishing a structure to quantify the disparities between genes and/or cells, and the resulting internal conflicts these cause within organisms.

### Can internal conflicts undo an organism?

Dawkins’ Paradox of the Organism asks why organisms aren’t torn apart by their selfish replicators. Looked at another way, the Paradox asks why the major transitions in individuality do not seem to go in reverse. Organisms are collectives made of parts (e.g., genes, genomes, and cells) that had had their own purposes earlier in evolutionary history but that can now function only as part of higher levels of individuality (West *et al*., 2015; Bourrat *et al*., 2022; Bourke, 2023; Doulcier *et al*., 2023). These ‘collective agents’ clearly harbor internal conflicts over the direction of the organism, but why do they not get out of hand? Or, in other words, why do we not observe “major reversions” where organisms are torn apart from within?

Our measures of unity provide a way to think about this question. One reason an organism might cease to operate as a collective is that its parts have such widely divergent goals for what the collective organism ought to be like. Our measure of trait disunity, *U*_*z*_, would capture this difference of optima. An analogy here might be helpful. A political party typically pursues the goals laid out in the party program. Not all party members, however, will agree with all of these goals, and would prefer the party to pursue alternative ones. These members may use different strategies to change the course of action taken by the party. They may, for example, seek out higher positions within the party to replace members from a different faction or they may try to convince members to vote for their proposed policy changes, just like how selfish elements may promote their own transmission and/or distort organismal traits for their own good (Patten *et al*., 2023). Successful parties tend to be coalitions of different factions, and a party whose members hold only slightly different goals for the party as a whole may reach a consensus that is satisfactory to all party members. However, parties where members hold more severe disagreements may be unable to do so. Instead, its members may split off and continue as independent politicians, or a schism in the party may yield two or more separate parties, each of which pursues a distinct party program. Our measure of trait unity would capture this difference and would point to the latter party as more likely to undergo dissolution.

Another reason that organisms might cease to function as a collective is that the fitness effects of the internal conflicts are simply so severe. Our fitness unity metric, *U*_*w*_, captures the potential for a loss of organismal fitness. Where this alteration is most severe, one would predict the greater risk for a major reversion in individuality. That said, these sorts of potential alterations are also the most likely to be thwarted by adaptations of the organism to suppress selfishness (Scott & West, 2019).

While many conceptual issues remain, our work provides several key insights pertaining to internal conflicts and their impact on organismal unity of purpose. First, internal conflicts can challenge organismal unity and may separately affect trait and fitness unity (Eqs. 3a and 3b). These two forms of unity are not equivalent (Figure 1). Second, different interpretations of the framework allow for different approaches to modelling internal conflicts. This includes different definitions of the organism and its evolutionary optimum 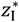, as well as its constituent genic factions and their involvement *p*_*i*_ (i.e. share, power, or potential; Figure 4). Third and final, the assumption that organisms are unified agents of evolution is often violated. An organism is not the sum of its parts. Instead, organisms are adaptive compromises – collectives in need of unification in their pursuit of fitness maximization. Taken together, our quantification provides a first formalization of the Paradox of the Organism, thus establishing a novel framework for understanding the evolution of organismality.

## Author contributions

Conceptualization: M.M.P. and J.A.Å.; funding acquisition: M.M.P. and J.A.Å.; model development: M.A.S.; simulations: M.A.S.; data analysis: M.A.S.; writing (original draft): M.A.S.; writing (review and editing): M.A.S., M.M.P. and J.A.Å.

## Acknowledgements

We thank the “Modeling Agency Formally” cluster for feedback on an earlier version of this manuscript.

## Data and code availability

Model source code, primary data, analysis scripts, and all resulting outputs are freely available via GitHub (https://github.com/MartijnSchenkel/InternalConflictQuantification).

## Supplementary Methods

### Genomic imprinting

Genomic imprinting is the process whereby the expression of an allele depends on whether it was inherited from the mother or from the father. The kinship theory (Haig, 2000) posits that genomic imprinting involves because of the matrigenic and patrigenic genome experience different relatedness to the bearer’s kin. That is, when for example a female mates with one male to produce the focal offspring, and with another male to produce another offspring, then the matrigenic genome has some probability of being represented in this second offspring. The patrigenic genome from the first offspring however has no representation in the second offspring (similarly, the patrigenic genome from the second offspring is not represented in the first). The inclusive fitness value of this second offspring thus differs from the perspective of the matrigenic and patrigenic genome halves in the focal offspring. A key consequence is then that social behaviors that affect the (inclusive) fitness of the focal offspring are differently appreciated by both genome halves as well.

We capture this internal conflict in a model where maternal resource investment into offspring survival is considered from the perspective of the organism, as well as its matrigenic and patrigenic genome halves. Each party has a different appreciation of the inclusive fitness value of the mother’s further reproductive success, measured in the production of additional siblings. In our model, mating is random, and each mating interaction yields a single offspring. Females mate multiply to generate multiple offspring and we assume that females remate exclusively with other males so that her offspring are all maternal half-siblings, but paternally unrelated. Each mother starts with a certain amount of resources, *R*. Producing offspring requires an investment of this maternal resource, the amount of which we assume is under the control of the offspring. The resource level obtained by offspring, *z*_I_, determines the probability of offspring survival *S*(*z*_I_). We are interested in determining the optimal outcome from the perspective of this offspring, and the role of internal conflict between its matrigenic and patrigenic genome, which represent the two genetic factions (*k* = 2) in this model. We use *i* ∈ {mat, pat} to refer to the matrigenic and patrigenic genome sets.

Offspring maximize their inclusive fitness by weighing their own probability of survival, and the survival of any kin produced by this mother:

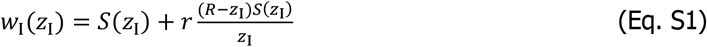

Here, *r* denotes the relatedness between the focal offspring and its kin 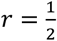 for full sibs, 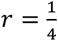 for half-sibs). The first term represents the expected survival of the focal offspring (direct fitness), whereas the second term represents the expected cumulative survival of its kin (indirect fitness). As we assume full remating, all offspring produced by a given female are maternal half-siblings of each other. We nonetheless assume that the investment level *z*_I_ is identical across all offspring produced by a female, which implies selection acts efficiently relative to the mutation rate so that beneficial mutations are rapidly fixed.

When females mate multiply, offspring relatedness drops, but this affects matrigenic and patrigenic genes differently. Patrigenes in a focal offspring have no future representation in kin produced by the same mother (*r* = 0). Thus, patrigenes optimize their own future representation by maximizing the likelihood of survival by the offspring so that:

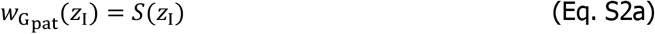

Conversely, matrigenes retain the possibility of being inherited through siblings to the offspring and therefore continue to include this when optimizing their inclusive fitness (Eq. S1, with 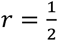):

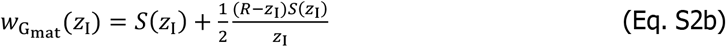

We implement the following survival function, adapted from Macnair & Parker (1978); Parker & Macnair (1978, 1979):

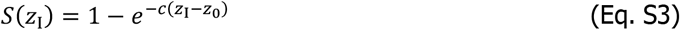

Here, *c* is a scaling parameter that determines how rapidly offspring survival increases with additional expenditure (higher *z*_I_), and *z*_0_ indicates the minimal investment for offspring survival to exceed 0. To show how investment affects the fitness of various genetic factions, we take the derivatives of offspring, maternal, and paternal fitness with respect to *z*_I_, yielding the following:

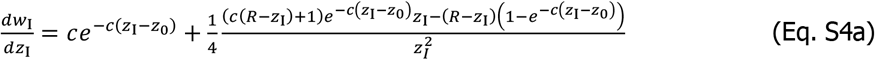

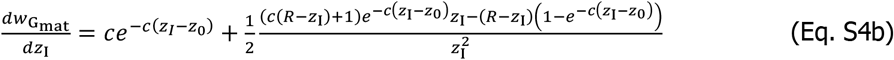

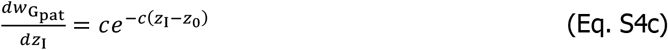

In Eq. S4a-c, the first term represents the fitness effect of increasing investment *z*_I_ on the survival of the offspring itself, whereas the second term in Eq. S4a,b represents the fitness effect on cumulative kin survival; this term is ignored for the patrigenic genome set as it has no representation in the offspring’s maternal half-siblings. To determine the genic-optimal values we solve 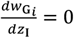, and for individual fitness we solve 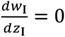 via numeric approximation (see “Model implementation and data analysis” below). We determine trait and fitness unities (*U*_*w*_ and *U*_*z*_) as described above. To determine *U*_*z*_, we assume that *z*_I_ ∈ [*z*_0_, *R*].

### Sex ratio distortion

Genomic elements such as sex chromosomes and cytoplasmic elements are transmitted to and from the two sexes in different ways. These asymmetrical segregation patterns lead genomic elements to differently value females and males (Cosmides & Tooby, 1981), as they are transmitted by the sexes at different rates and therefore can spread more easily through one sex than through the other. Following from this, genomic elements may disagree over the optimal sex ratio among an individual’s offspring. We capture this internal conflict by determining these differences play out in the context of local mate competition (Hamilton, 1967).

Under local mate competition theory, female foundresses establish demes of size *n*. They produce a number of offspring *o*, with a sex ratio *z*_I_, so that the number of (expected) sons equals *z*_I_*o* and daughters (1 − *z*_I_)*o*. For simplicity, we assume *o* to be constant so that it can be ignored in any further analysis. Daughters and sons of all foundresses mate and compete for mating in the natal deme, after which the inseminated daughters disperse to occupy novel demes; sons die in their natal deme. This population structure promotes a female-biased sex ratios 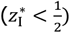 at low deme sizes, which increases towards equality 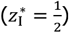 as *n* increases (see Main text). As described above, specific genetic elements have different optimal sex ratios, which similarly vary based on deme size.

Our model of internal conflict over optimal sex ratio aims to capture the severity of this conflict in relation to the deme size, and the relative share *p*_*i*_ of genomic elements that exhibit different optimal sex ratios 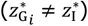. We first consider the fitness of foundresses and how it is shaped by the sex ratio of offspring they effect. In a deme with *n* − 1 foundresses all producing the resident sex ratio 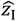, a mutant foundress producing a sex ratio *z*_I_ will have fitness *w*_I_ given by:

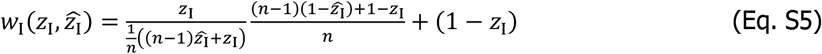

Here, the first term represents the number of fertilizations obtained by her sons, and the second represents the production of daughters. The optimal sex ratio 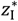 is then given by:

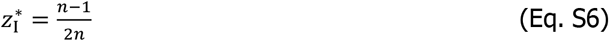

This optimal sex ratio is the resident strategy 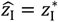 for which all other sex ratio strategies 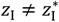 yield a lower fitness than for 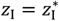, i.e. for which all mutants have reduced fitness compared to residents. Given a resident strategy 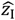, the derivative of Eq. S5 with respect to the focal individual’s strategy *z*_I_ is given by:

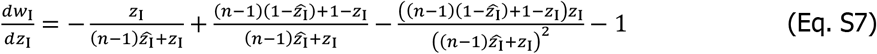

A caveat is that various genomic elements, such as sex chromosomes, which are not represented equally in the two sexes, are not transmitted equally to and from daughters and sons. While autosomes segregate equally, the sex chromosomes (X vs. Y, Z vs. W) segregate differently to and from males and females, and similarly cytoplasmic elements are predominantly inherited maternally (Birky, 2001). This results in different optimal sex ratios for these elements, which in case of the major sex chromosomes (X- and Z-chromosomes) additionally depends on whether they occur in males or females (and further assumes that males can exert control over the sex ratio of their brood). Hamilton (1979) showed that these optima correspond to:

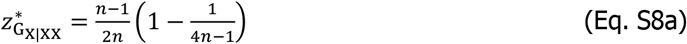

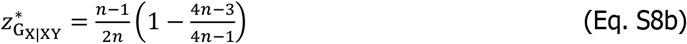

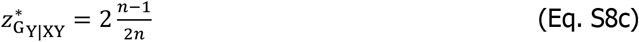

Here, 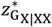 refers to the optimal sex ratio for an X-chromosome in an XX female, and 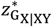 and 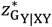 gives the optimal sex ratios for X- and Y-chromosomes in XY males. Similarly, for Z- and W-chromosomes in ZZ males and ZW females:

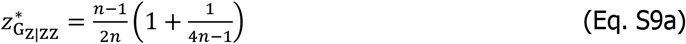

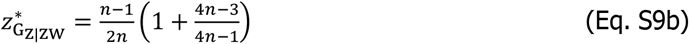

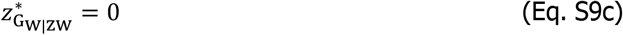

Note that the strategy for the W-chromosome is identical to that of cytoplasmic elements under strict maternal inheritance, as both exhibit the same female-limited segregation pattern. When patches contain only a single foundress, all genic optima correspond to *z*_I_ = 0, equaling the individual-optimal value. In practice, an optimal sex ratio of 0 means individuals should produce the minimal number of sons that would be capable of fertilizing the comparatively vast number of daughters. When patches are infinitely large, i.e. when the population is effectively a single large and panmictic patch, the organismal optimum corresponds to equal sex ratio 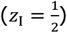, whereas the optimal values for the Y- and W-chromosomes are 1 and 0, respectively, representing all-or-nothing strategies.

In analyzing our model, we assume that all resident foundresses in a patch use the individual-optimal strategy, i.e. 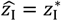. For the focal (i.e. mutant) foundress, we determine *U*_*w*_ and *U*_*z*_, by assuming she plays a mutant strategy *z*_I_ that is given by the different genic optima. This situation is used to determine how severe internal conflict is under the assumption that individuals are optimized. We assume the sex chromosomes (XX and XY, ZZ and ZW) represent a fraction *p* of the genome, and the remainder is represented by the autosomes. In XY (ZW) individuals, we assume the X- and Y-chromosomes (Z- and W-chromosomes) represent an equal fraction of *p*, i.e. *p*_X_ = *p*_Y_ = *p*/2. For simplicity, we ignore the contribution of cytoplasmic elements, who would like W-chromosome have an optimal sex ratio 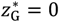. Note that the optimal strategies of the autosomes, and the X-chromosomes in XX females, assume maternal control over the sex ratio, whereas for the X- and Y-chromosome in males, paternal control is implied.

### Model implementation and data analysis

The genomic imprinting and sex ratio models were implemented in C++. For the genomic imprinting model, we determined the evolutionary optima of each entity (organism, maternal genome, paternal genome) by approximating the real-valued roots of the derivatives for the fitness of individuals 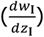 and of the maternal and paternal genome halves 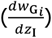 using a bisection algorithm. This was used to approximate the derivative up to an accuracy ε = 1 × 10^−10^, provided that a root could be found. The fitness function for the patrigenic genome is monotonically increasing, and hence its derivative has no roots. The fitness optimum is instead equal to the cumulative amount of resources *R* of the mother, as this value optimizes fitness under the current constraints.

For the sex ratio model, we evaluated the fitness of individuals playing a strategy *z*_I_ that equals either the individual-optimal strategy 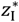 or a strategy 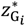 that is optimal for one of its genetic factions *i*. We assumed that all other foundresses in the deme play the individual-optimal strategy 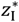. We further assumed that the sex chromosomes (XX in females, XY in males) represent a combined share *p* of the genome, and that in XY males the X and Y both represent a share *p*_X_ = *p*_Y_ = *p*/2; the autosomes, which follow the same optimization program as the individual as a whole, represent a share 1 − *p*. For simplicity, we ignore the cytoplasmic elements, whose optimal strategy is to produce only females 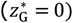. Because our analysis revealed that individuals in female heterogametic systems yield identical unity scores as those in male heterogametic systems (but with sexes reversed, i.e. XY males comparable to ZW males, XX females comparable to ZZ males), we restrict our discussion in the main text to XX females and XY males.

All data was analyzed and visualized in R (v. 4.0.2; R Development Core Team, 2023) and RStudio (v. 2022.07.01; RStudio Team, 2023) using the ‘cowplot’, ‘tidyverse’, and ‘viridis’ packages (Garnier, 2018; Wickham *et al*., 2019; Wilke, 2019).

## Supplementary Figures

**Supplementary Figure 1:**
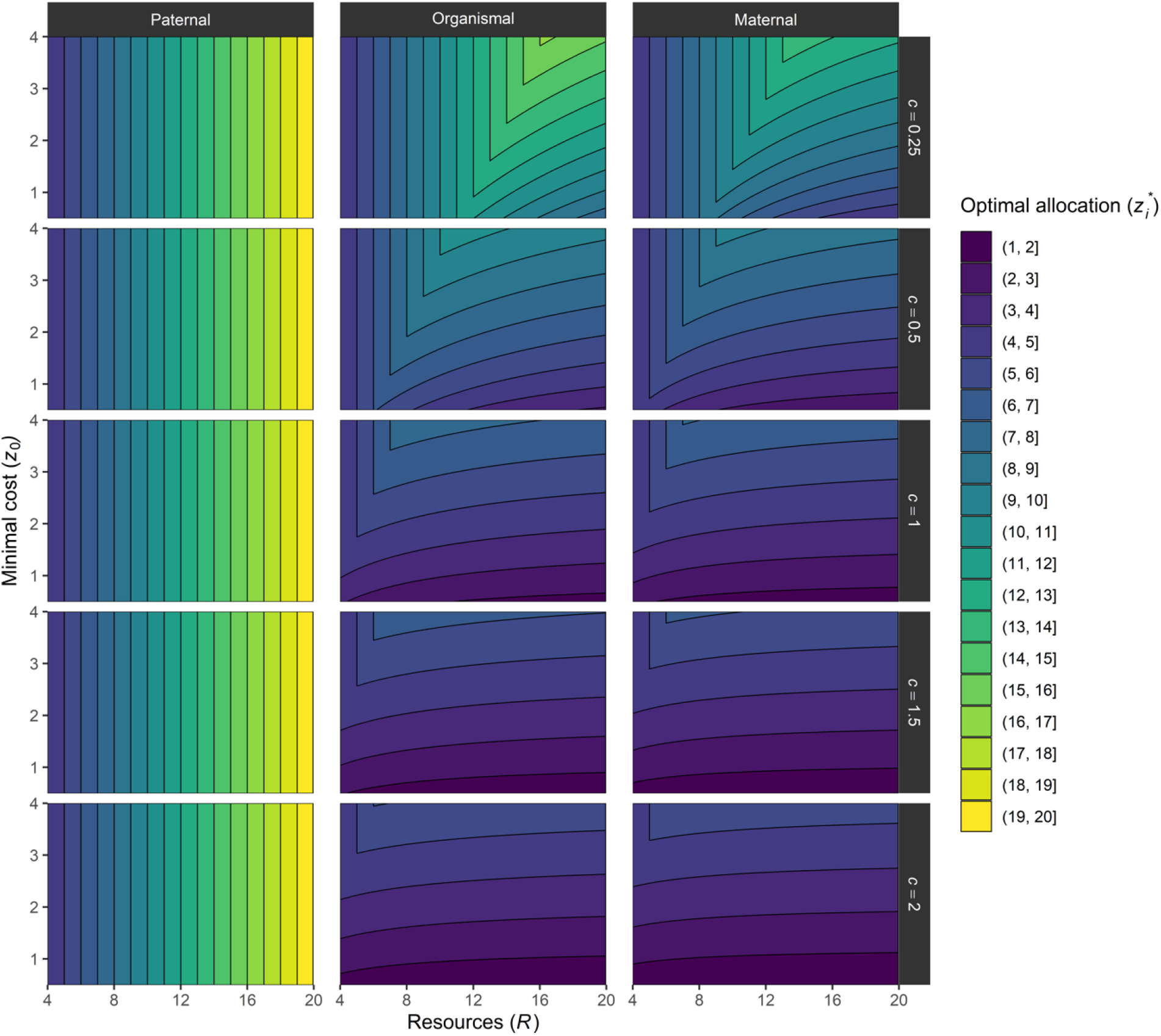
Optimal resource allocation strategies 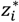 of the organism 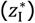, the patrigenic genome 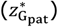, and the matrigenic genome 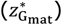 under the genomic imprinting model. Mothers mate multiply, so that all siblings are half-siblings via the offspring’s mother. Relative to the focal offspring, its matrigenes overvalue the indirect fitness value of these siblings, favoring a reduced investment into the focal offspring. Patrigenes are not represented in any of the focal offspring’s siblings, so that they experience no indirect fitness through them and thus value overinvestment into the focal offspring.

**Supplementary Figure 2:**
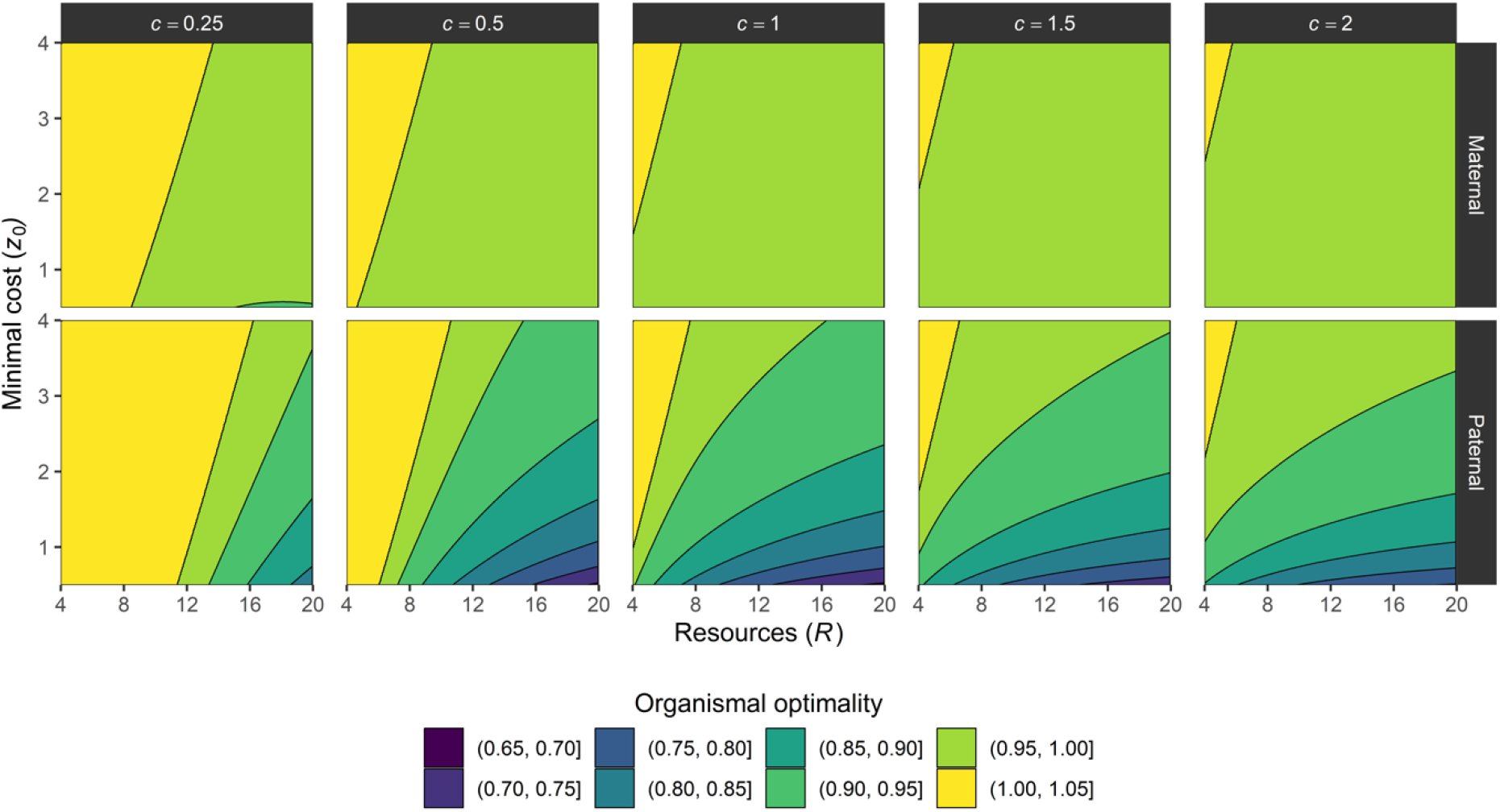
Organismal optimality under genomic imprinting when maternal investment is under the control of the matrigenic (top row) or patrigenic genome (bottom row). Optimality is defined as the fitness of the individual at the matrigenic/patrigenic genome optimum 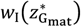 or 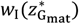 divided by its fitness at the individual optimum 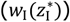.

**Supplementary Figure 3:**
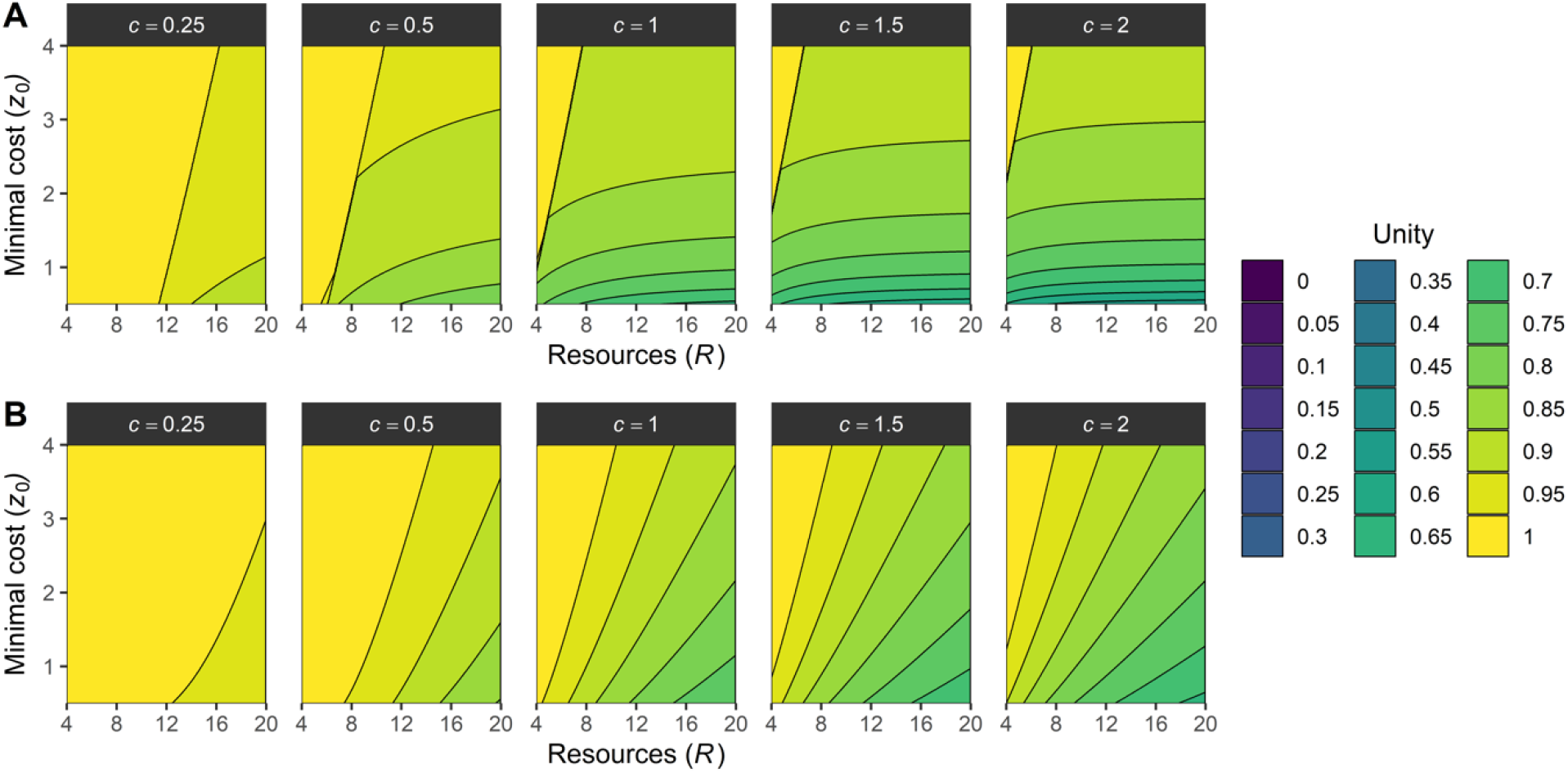
(A) Trait unity (*U*_*z*_) and (B) fitness unity (*U*_*w*_) of parental investment in light of genomic imprinting. Optimal parental investment is investigated from the perspective of their offspring; increased investment enhances their survival, but reduces the production of siblings. Owing to relatedness differences, matrigenes and patrigenes favor under- and overinvestment of resources relative to the individual-level optimum, reducing the trait unity. In particular, patrigenes favor complete investment of all maternal resources *R* as they have no representation in any of the other offspring produced. This difference drives reduced trait and fitness unity as it harms the indirect fitness component of the focal offspring’s inclusive fitness. *z*_0_ represents the minimal investment, whereas *c* represents how rapidly survival increases with further investment. Lower minimal investment (*z*_0_) and/or higher efficiency (*c*) leads to even stronger reductions in trait and fitness unity through the patrigenes’ preference for overinvestment.

**Supplementary Figure 4:**
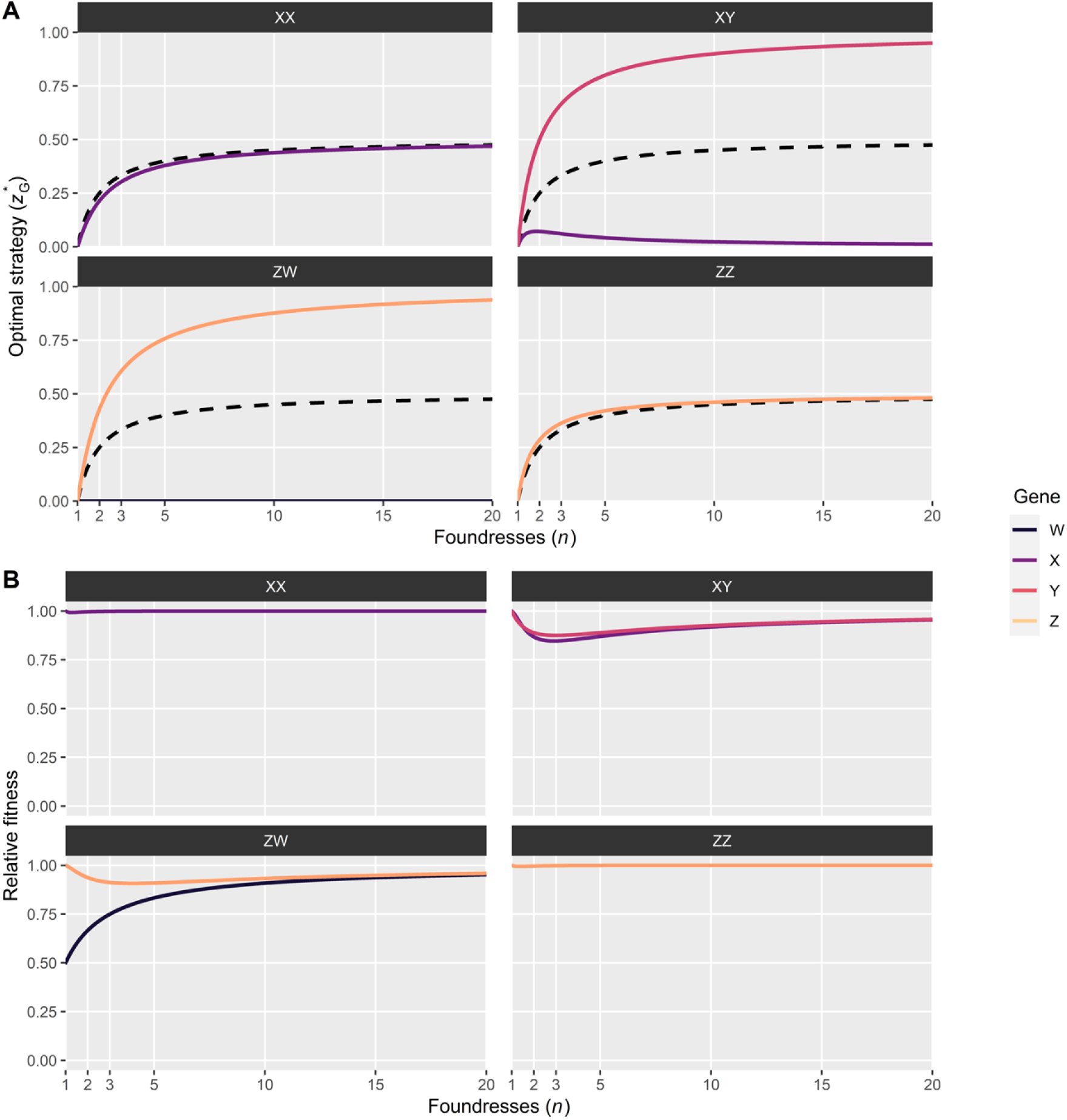
Optimal sex ratios under local mate competition of the organism, its X-chromosome, and its Y-chromosome (or Z- and W-chromosome in ZZ/ZW individuals). Autosomal optima are equivalent to organismal optima and are therefore not depicted. (A) Optimal strategies 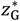 for X- and Y-chromosomes in XX and XY individuals, or Z- and W-chromosomes in ZZ and ZW individuals; the dashed black line represents the individual-optimal strategy 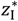. (B) Fitness of the individual under the optimal strategies 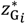 of its sex-chromosomes relative to its fitness under the individual-optimal strategy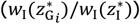. In determining fitness, we assume the remaining *n* - 1 foundresses in the focal individual’s patch use the individual-optimal strategy 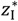, and that the focal individual plays the strategy that is optimal to the specific sex chromosomes as depicted in (A).

**Supplementary Figure 5:**
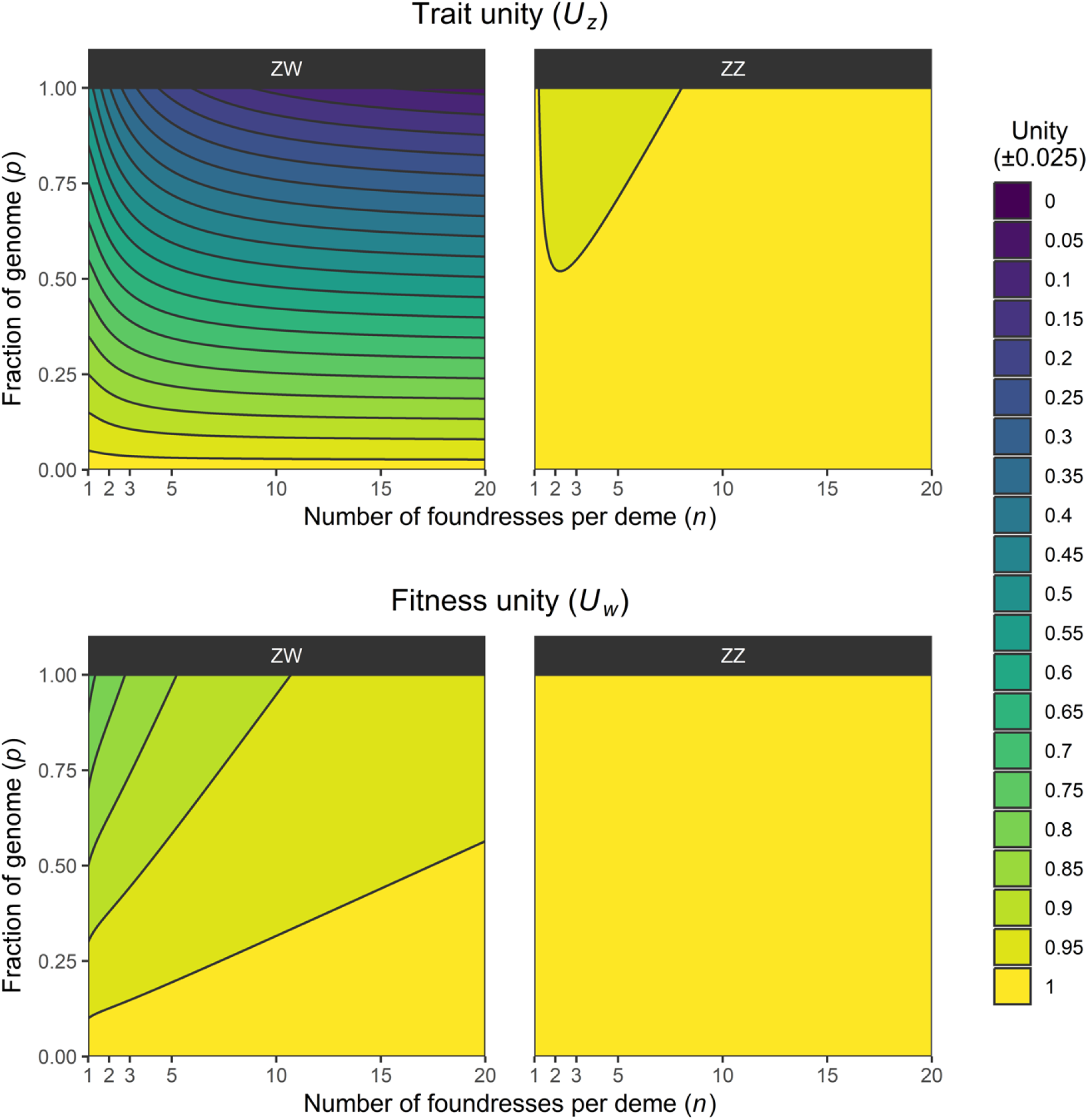
Trait unity (top) and fitness unity (bottom) with regard to sex ratios under local mate competition in ZW versus ZZ individuals. In determining trait and fitness unity, all *n* − 1 residents in the patch play the individual-optimal strategy 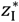. The focal individual consists of a genome of which a proportion 1 − *p* is autosomal, and therefore has the same optimal strategy as the individual. The remainder *p* is distributed among the Z- and W-chromosomes, which have genic-optimal strategies 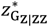 (in ZZ individuals), 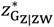, or 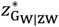 (in ZW individuals); in ZW individuals, the Z- and W-chromosome represent an equal portion *p*/2 of the genome.

